# Immunoprofiling reveals novel mast cell receptors and a continuous nature of human lung mast cell heterogeneity

**DOI:** 10.1101/2021.03.12.435093

**Authors:** Elin Rönnberg, Daryl Zhong Hao Boey, Avinash Ravindran, Jesper Säfholm, Ann-Charlotte Orre, Mamdoh Al-Ameri, Mikael Adner, Sven-Erik Dahlén, Joakim S. Dahlin, Gunnar Nilsson

## Abstract

**Background:** Immunohistochemical analysis of granule-associated proteases have revealed that human lungs mast cells constitute a heterogeneous population of cells, with distinct subpopulations identified. However, a systematic and comprehensive analysis of cell surface markers to study human lung mast cell heterogeneity is yet to be performed.

**Methods:** Human lung mast cells were obtained from lung lobectomies and the expression of 332 cell surface markers were analyzed using flow cytometry and the LEGENDScreen^™^ kit. Markers that exhibited a high variance were selected for additional analyses to reveal whether they correlated and if discrete mast cell subpopulations were discernable.

**Results:** We identified expression of 102 surface markers on human lung mast cells. Several markers showed a high continuous variation of expression within the mast cell population. Six of these markers correlated: SUSD2, CD49a, CD326, CD34, CD66 and HLA-DR. The expression of these markers also correlated to the size and granularity of the mast cells. However, no marker produced an expression profile consistent with a bi- or multimodal distribution.

**Conclusions:** LEGENDScreen analysis identified more than 100 cell surface markers on mast cells, out of which 23 have to our knowledge not previously described on human mast cells. Several of the newly described markers are known to be involved in sensing the microenvironment, and their identification can shed new light on mast cell functions. The exhaustive expression profiling of the 332 surface markers failed to detect distinct mast cell subpopulations. Instead, we demonstrate a continuous nature of human lung mast cell heterogeneity.

## Introduction

Heterogeneity among mast cells has been known for long time and was first attributed to differential expression of proteoglycans in rodent mast cells, which gave them distinct staining patterns ^1^. This led to the division of rodent mast cells into connective tissue mast cells and mucosal mast cells. In humans, mast cell heterogeneity has been based on the expression of mast cell proteases, i.e., the expression of tryptase only (MC_T_) or those expressing both tryptase and chymase (MC_TC_) as well as carboxypeptidase A ^2,3^. These subtypes have been defined using immunohistochemistry, a method that produced binary results, that is absence or presence of expression. The MC_TC_ subtype is more predominant in connective tissues such as in the skin, while the MC_T_ subset is more prevalent in mucosal surfaces such as in the airways and the gastrointestinal tract ^4^.

Mast cells are found in the human lung in all different compartments; i.e., under the epithelium, in smooth muscle bundles, around pulmonary vessels, in the parenchyma and in close proximity to sensory nerves ^5^. Human lung mast cells (HLMC) are attributed to several important functions in health and diseases, such as host defense, induction of acute inflammatory responses, vascular regulation, bronchoconstriction and tissue remodeling ^6–9^. Heterogeneity of HLMCs was first described to be related to differences in cell size and functionality; i.e., response to secretagogoues^10,11^. Later it was described that the MC_T_ subtype is the predominant subtype in the lung, except around pulmonary vessels where MCT and MC_TC_ are found in equal numbers ^2^. However, the heterogeneity among HLMC goes beyond size and protease expression as demonstrated by the differential expression of certain mast cell related markers (FcεRI, IL-9R, 5-LO, LTC_4_S etc.) among MC_T_ and MC_TC_ populations in different lung compartments ^12^.

Mast cell heterogeneity have primarily been studied in a binary manner using immunohistochemistry, describing absence or presence of expression. Here, we used a quantitative flow cytometry-based approach to study HLMC heterogeneity, profiling the expression of 332 markers. None of these markers distinctly divided the mast cells into subpopulations. However, several markers showed a high degree of variation within the mast cell population with gradient-non-clustered expression pattern. Six of these markers correlated to each other, revealing a continuous nature of HLMC heterogeneity rather than distinct subpopulations.

## Materials and Methods

### Ethical Approval

The local ethics committee approved the collection of lung tissue from patients undergoing lobectomies, and all patients provided informed consent (Regionala Etikprövningsnämnden Stockholm, 2010/181-31/2).

### Cell preparation

Single cell suspensions from macroscopically healthy human lung tissue were obtained as previously described ^13^. Briefly, human lung tissue was cut into small pieces and enzymatically digested for 45 min with DNase I and Collagenase. Thereafter, the tissue was mechanically disrupted by plunging through a syringe, the cells were washed, and debris was removed by 30% Percoll centrifugation. The cells were, after preparation, stained and analyzed by flow cytometry.

### Flow cytometry

The following antibodies were used for surface staining: CD45-V500 (Clone HI30, BD Biosciences, San Jose, CA, USA), CD14-APC-Cy7 (Clone M5E2, Biolegend, San Diego, CA, USA), CD117-APC (clone 104D2, BD Biosciences), FcεRI-FITC (clone CRA1, Miltenyi Biotec, Bergisch Gladbach, Germany), FceRI-PE (clone CRA1, Biolegend), SUSD2-PE (clone W3D5, Biolegend), CD63-PE/Cy7 (Clone H5C6, BD Biosciences), CD49a-BV786 (Clone SR84, BD Biosciences), CD66a/c/e-A488 (clone ASL-32, Biolegend), CD326-BV650 (Clone 9C4, Biolegend), CD34-BV421 (Clone 581, BD Biosciences), HLA-DR-PE/Cy5 (Clone L243, Biolegend), CD344-PE/Vio770 (Clone CH3A4A7, Miltenyi Biotec). For the LEGENDScreen^™^ human cell Screening kit, that contain 342 antibodies all conjugated to PE (Cat. 70001, Biolegend), detailed in Supplementary Table S1, cells were first stained with CD45, CD117, CD14 and FceRI antibodies, thereafter they were stained with the kit according to the manufacturer’s instructions. Mast cells were gated as CD45^+^, CD14^low^, CD117^high^ (Figure 1). For intracellular staining, cells were fixed with 4% PFA and permeabilized using PBS-S buffer (0.1% saponin in PBS with 0.01 M HEPES). Unspecific binding was blocked using blocking buffer (PBS-S with 5% dry milk and 2% FCS). The cells were thereafter stained with tryptase antibodies (clone G3, Millipore, Burlington, MA, USA) conjugated in-house with an Alexa Flour 647 Monoclonal antibody labeling kit (Thermo

**Figure 1.**
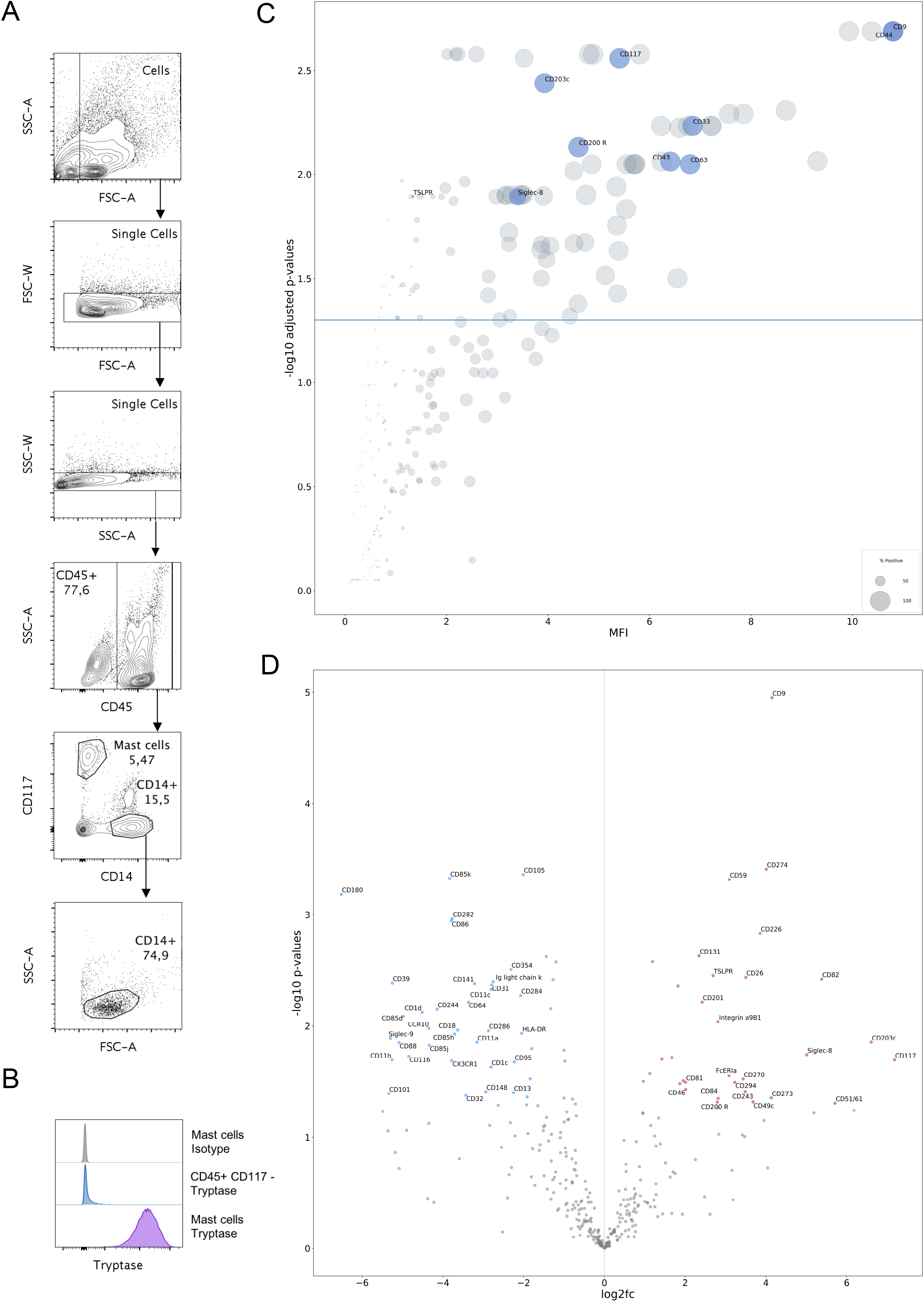
Gating strategy and LEGENDSscreen results. Single cell suspensions of human lung tissue were stained with CD45, CD114, CD14 and FcεRI and thereafter the LEGENDScreen human cell Screening kit. (A) A representative of the gating strategy of human lung mast cells and CD14^+^ cells are shown. (B) Intracellular tryptase stained human lung mast cells, compared to isotype stained mast cells and tryptase stained CD45^+^, CD117^-^ cells. (C) Scatter plot of p-values, MFI and percent positive of each marker on the human lung mast cells. Y-axis plots −log10 FDR-adjusted p-values from 2-sided individual t-tests (marker against FMO controls), blue line represents the confidence cut-off of −log10 (0.05). X-axis plots normalised ln(MFI) values (marker subtracted by plate matched FMO control) and size of circles represents percentage positive cells with the positive gate set according to the FMO. Some mast cell markers are highlighted in blue. (D) Comparison of marker expression on mast cells and CD14^+^ cells. Volcano plot showing log2-fold change of mast cells divided by CD14^+^ cells (normalized MFI values with plate matched FMO subtracted) against −log-10 p-values (independent 2-sided t-test) of mast cells against CD14^+^ cells. Markers are only annotated if abs(log2fc) => 2, and p-value < 0.05. n=3.

Fisher Scientific, Waltham, MA, USA), or CPA3 antibodies (clone CA5, a kind gift from Andrew Walls, Southampton, UK) conjugated in-house with an Alexa Fluor^™^ 488 Antibody Labeling Kit (Thermo Fisher Scientific). The cells were analyzed using a BD FACSCanto (BD, Franklin Lakes, NJ, USA) or BD LSRFortessa, and FlowJo software version 10 (FlowJo LLC, Ashland, OR, USA) was used for flow cytometry data analysis.

### Statistics

Statistical analyses were performed with GraphPad Prism software version 7.0b, or the Python environment (3.7) with the following packages: statsmodels (0.10.1), seaborn (0.9.0), scipy (1.4.1), pandas (1.1.0), numpy (1.18.1), matplotlib (3.1.3). For specific methods used see figure legends. * p < 0.05; ** p < 0.01; *** p<0.001; **** p<0.0001.

## Results

### Immunoprofiling of human lung mast cells

The expression of cell surface antigens was thoroughly investigated by flow cytometry using the LEGENDscreen^™^ kit containing 342 antibodies, including 10 isotype controls. The HLMCs were gated as CD45^+^, CD14^low^, CD117^high^ (Figure 1A) and the gated HLMC population expressed high levels of tryptase, confirming the identity of the gated cells (Figure 1B). The expression of some relevant mast cell markers included in the LEGENDscreen are highlighted in figure 1C showing the percent positive cells, the median fluorescent intensity (MFI) and the significance of expression within the HLMC population. Many of the highly expressed markers on HLMCs are broadly expressed, such as β2-microglobulin (B2M), CD44, and CD9 (Figure 1C). To determine which of the markers are more relevant for HLMCs, we compared the expression to CD45^+^CD14^+^ CD117^-^FSC^int^SSC^low^ cells (Figure 1A). Well-known monocyte markers such as CD11b, CD11c, CD31, CD141, CXCR1 and HLA-DR had higher expression on the CD14^+^ cells, whereas classical mast cell markers such as CD117, FceRI, CD203c, Siglec-8, and TSLPR, showed a higher expression on the HLMCs. The highest significant differences of HLMCs to the CD14+ cells included; CD9, CD59, CD274 and CD226 (Figure 1D). CD9 is a broadly expressed tetraspanin with a wide variety of functions, in mast cells it is abundantly expressed and has been implicated in chemotaxis and activation^14^. CD59 can prevent complement induced cytolytic cell death by preventing assembly of the complement membrane attack complex and have also been implicated in T cell activation^15^. CD274 is also known as programmed death ligand-1 (PD-L1) and can cause blockade of T-cell activation^16^. CD226 has received increasing interest in recent years and can play a role in many immunological processes^17^ including enhancement of FcεRI mediated activation in mast cells^18^. HLMCs significantly expressed (MFI compared to the fluorescent minus one (FMO) control) 102 out of the 332 markers included in the LEGENDScreen (Figure 2). Surface expression of 23 of those have to the best of our knowledge not been described on (non-neoplastic) human mast cells before (Table 1).

**Figure 2.**
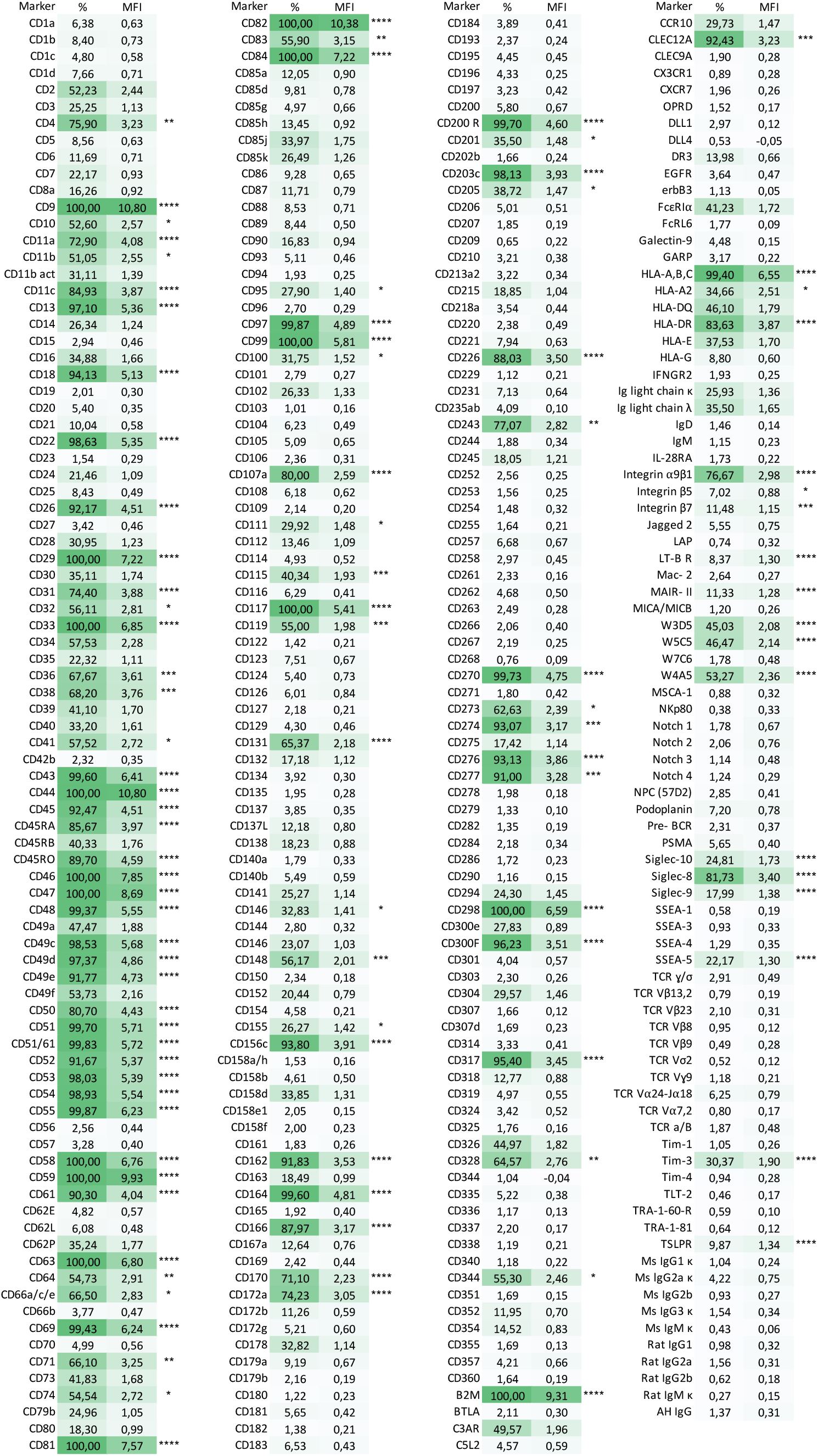
Expression of cell-surface antigens of human lung mast cells. Single cell suspensions of human lungs were stained with CD45, CD117, CD14 and FcεRI antibodies and thereafter stained with the LEGENDScreen human cell screening kit. Mast cells were gated as CD45^+^, CD14^low^ and CD117^high^. Shown are the percent positive (%) for each marker and the MFI that was normalized to the plate matched FMO control and log10 transformed. Significance of the MFI compared to the FMO control is shown (one-way Anova with Dunnett’s multiple comparisons test), * p < 0.05; ** p < 0.01; *** p<0.001; **** p<0.0001. n=3.

**Table 1.** Novel antigens identified on human lung mast cells

| Marker (clone) | Description |
| --- | --- |
| CD36 | Receptor binding a broad range of lipids |
| **CD45RO** | Isoform of CD45 |
| **CD66a/c/e** | Adhesion molecules |
| **CD74** | Involved in MHC class II antigen processing and a receptor for macrophage migration inhibitory factor. |
| **CD111** | Adhesion molecule |
| **CD115** | Receptor for M-CSF and IL-34. |
| **CD131** | Common $\beta$ subunit of the IL-3, IL-5 and GM-CSF receptors |
| **CD143** | Metallopeptidase |
| **CD148** | Tyrosine phosphatase involved in signal transduction |
| **CD164** | Sialomucin involved in cell adhesion and proliferation |
| **CD166** | Glycoprotein involved in cell adhesion and migration |
| **CD205** | Endocytic receptor involved in antigen uptake and processing |
| **CD243** | Involved in transportation of molecules across cell membranes |
| **CD270** | Receptor for TNFSF14, BTLA, LTA and CD160 |
| **CD277** | Regulate T cell responses |
| **CD317** | Blocks the release of certain viruses from infected cells |
| **CD344 (Frizzled-4)** | Receptor for Wnt proteins and norrin |
| **CLEC12A/CD371** | C-type lectin-like receptor with immunoreceptor tyrosine-based inhibitory motif (ITIM) |
| **Integrin $\alpha 9\beta 1$** | Integrin mediating cell adhesion and migration |
| **SUSD2 (W3D5, W5C5)** | Potentially involved in cell adhesion as this transmembrane protein contains functional domains associated with adhesion molecules |
| **(W4A5)** | Antigen has yet to be described |
| **Siglec-9** | Lectin that binds sialic acid and has ITIMs |
| **SSEA-5** | A glycan |

### Heterogeneous expression of the high-affinity IgE receptor, FcεRI

The LEGENDScreen analysis failed to capture significant staining of the high-affinity IgE receptor, FcεRI (Figure 2). However, the use of the same antibody clone in the backbone staining panel likely explains this observation. To investigate this further we studied the expression of FcεRI on mast cells separately from additional donors. The expression of CD117 and FcεRI from four donors are shown in Figure 3A. Approximately one half were ~100% positive for the marker (Figure 3B). Furthermore, even in the 100% positive individuals the level of expression, i.e., MFI, varied considerably (Figure 3C).

**Figure 3.**
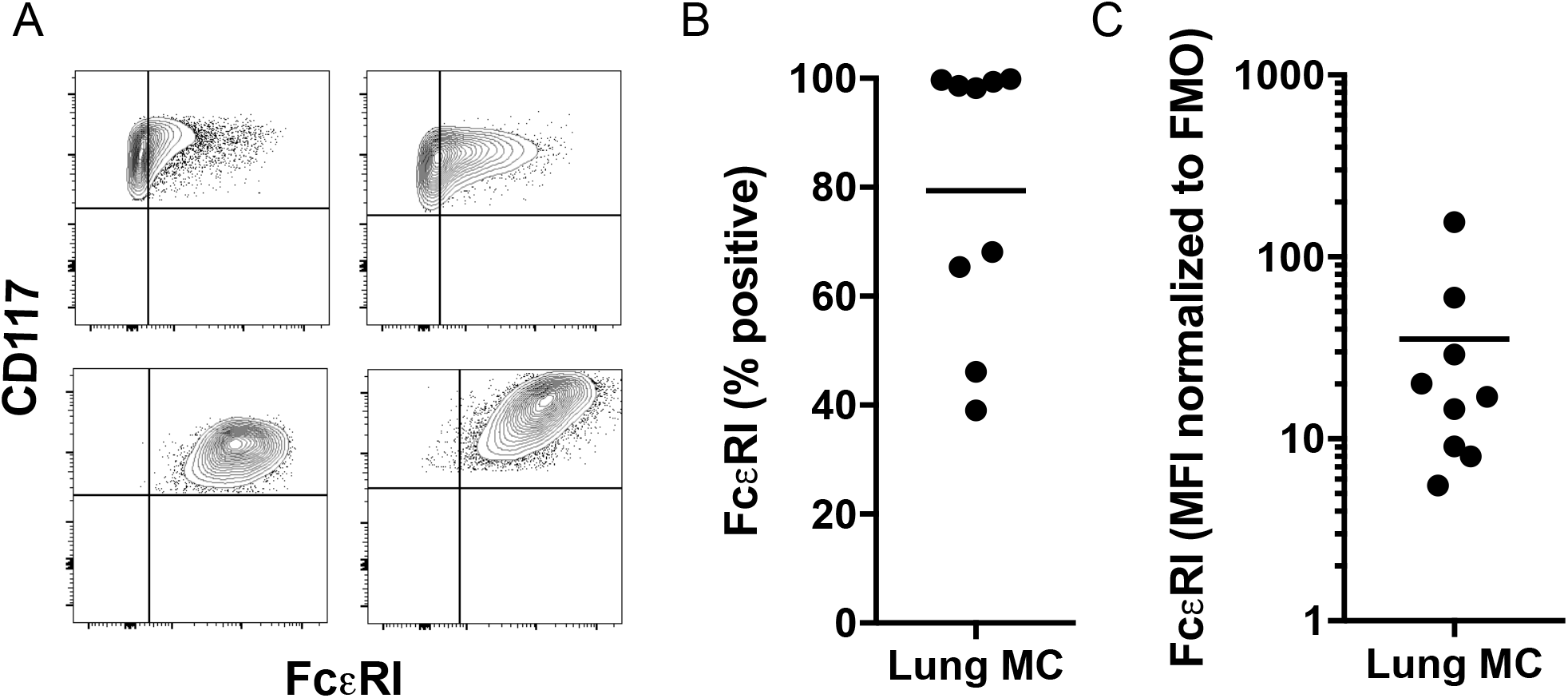
FcεRI expression on human lung mast cells. Examples of CD117/FcεRI expression on HLMCs gated as CD45^+^, CD14^low^ and CD117^high^ from four donors (A). Quantification of percent positive for FcεRI (B) and MFI of FcεRI normalized to the matched FMO control (C). n=9.

### Heterogeneous expression of cell surface markers with a continuous distribution

None of the markers clearly and consistently divided the HLMCs into subpopulations (data not shown). However, several markers showed a considerable continuous expression variation within the population, quantified by calculating the robust coefficient of variation (CV) (Table 2). The two antibodies with the highest CV was to the same antigen, SUSD2, a marker identified on mesenchymal and pluripotent stem cells with functional domains inherent to adhesion molecules ^19,20^. Co-stainings of the seven highest CV markers revealed that six of these markers correlated (SUSD2, CD49a, CD326, CD34, CD66 and HLA-DR), while CD344 did not correlate to any of the other markers (figure 4, FMO controls in supplementary figure S1). Furthermore, to investigate if these markers correlated with the classical mast cell subtypes, MCT or MCTC, co-staining with anti-CPA3 was performed but no correlation was observed (Figure 4G, FMO control in supplementary figure S1). In addition, these markers did not show co-staining with any of the other markers that were included for gating purposes in the LEGENDScreen, including CD45, CD14, CD117 and the FcεRI receptor (data not shown). Furthermore, cells high in SUSD2 showed a higher FSC and SSC, indicating that they are bigger and have a higher inner complexity, i.e, have more granules (Figure 4 H-J). SUSD2 have been linked to proliferation in cancer cells ^21^, why we investigated the proliferation status of the cells with the proliferation marker Ki-67. However, in agreement with that mast cells are long-lived cells with a low turnover ^22^ no staining was observed (Supplementary Figure S2.).

**Figure 4.**
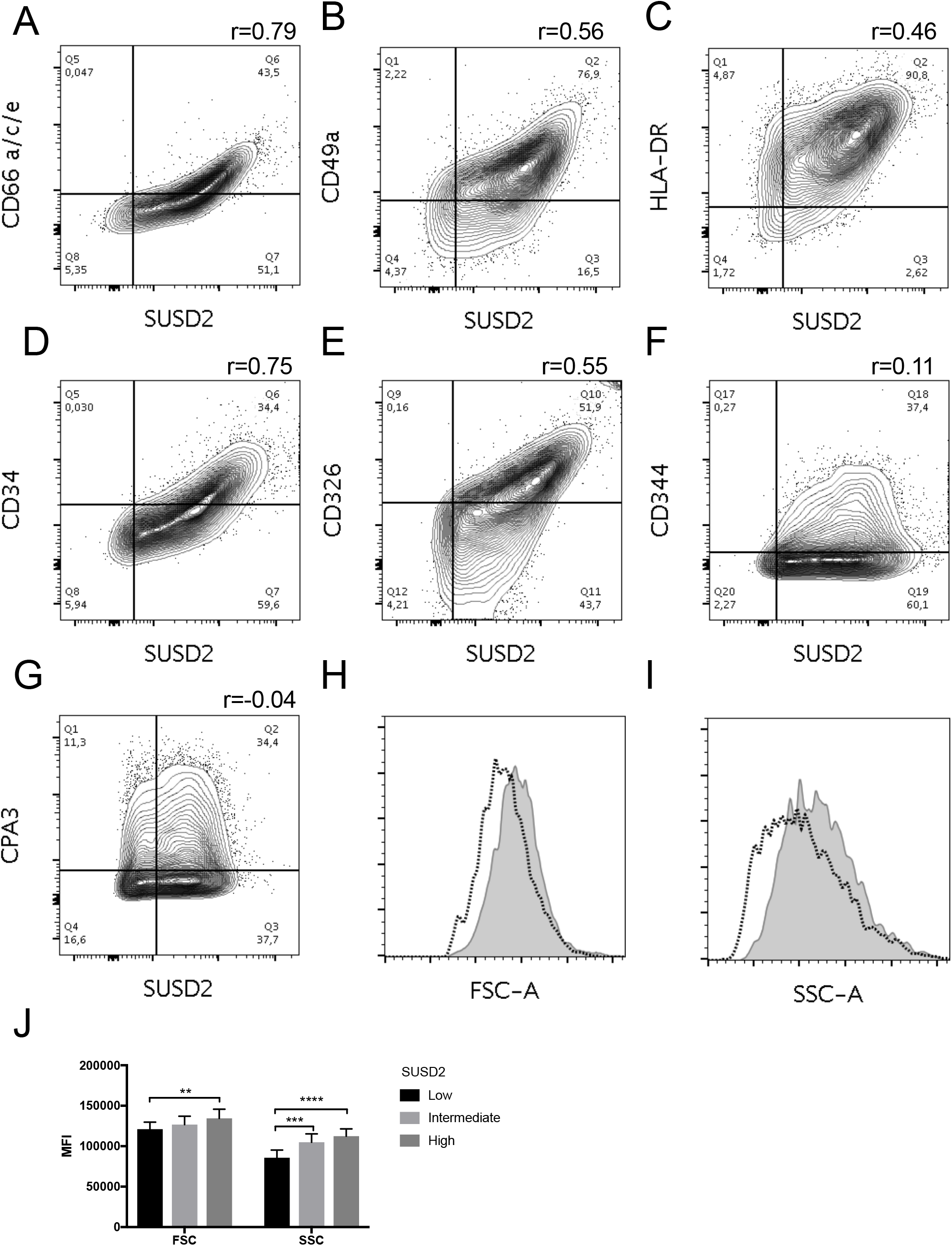
Correlations between markers with high CV, size and granularity. HLMCs gated as CD45^+^, CD14^low^ and CD117^high^ were co-stained with SUSD2 (A-H), CD66a/c/e (A), CD49a (B), HLA-DR (C), CD34 (D), CD326 (E), CD344 (F), and CPA3 (G). Representatives of 4 donors are shown. Pearson correlations to SUSD2 of the fluorescent intensity data in each donor was calculated using Graphpad prism and the average r value of four donors is shown. All correlations had a p>0.0001. SUSD2low, intermediate and high cells were gated and FSC (H) and SSC (I) of the low (dotted line) and high (filled grey) SUSD2 populations are shown. Quantification of FSC and SSC is shown in (J), mean ± SEM, n=5. Two-way Anova with Bonferroni’s multiple comparisons test was performed. * p < 0.05; ** p < 0.01; *** p<0.001; **** p<0.0001.

**Table 2.** The 10 markers from the LEGENDScreen with the highest Robust coefficient of variation (Robust CV)

| **Marker** | **Robust CV** |
| --- | --- |
| **SUSD2 (W5C5)** | 264 |
| **SUSD2 (W3D5)** | 246 |
| **CD344** | 172 |
| **CD49a** | 160 |
| **CD326** | 155 |
| **CD66a/c/e** | 153 |
| **CD34** | 134 |
| **HLA-DR** | 133 |
| **SSEA-5** | 131 |
| **CD63** | 124 |
| **CD38** | 123 |

## Discussion

Although attempts have been made to map cell-surface antigens on HLMCs ^23–28^, an extensive mapping including the heterogeneity of expression of the cell-surface antigens has not been carried out. In this study, we identified significant expression of 102 markers on the HLMCs out of which, to the best of our knowledge, 23 are novel mast cell markers (Table 1). Several of these markers are described as markers expressed on stem cells, including SSEA-5, SUSD2, W4A5, CD243, CD111, CD131 and CD164. The expression of stem cell markers on mast cells is in accordance with results from the FANTOM5 consortium, in which mast cells exhibit similarities with stem cells ^29^. In some cases, our results are in disagreement with previous published data, for example CD4, CD10, CD36 and CD74 have previously been shown to not be expressed by HLMC ^25,27^. This discrepancy might be explained by differences in the procedures, where to the contrary of published data we did not purify or culture the mast cells prior to analysis^24,26–28^. Culturing mast cells have been shown to alter their phenotype and expression of cell surface receptors before ^29,30^.

In an immunohistological study by Andersson et al. they found that the expression of the FcεRI receptor on HLMC differed within different compartments of the lung, with mast cells present in the parenchyma being negative for FcεRI ^12^. In our study, it was not a clear-cut division of a negative and positive FcεRI population but rather a continuous spectrum of different levels of expression and about 50 percent of the patients expressed FcεRI on virtually all of the mast cells (Figure 3A). These discrepancies could be due to different detection limits of the two different techniques used, immunohistochemistry and flow cytometry. We measured the expression in a quantitative manner using flow cytometry, thus finding that there is a spectrum of different expression levels, while in the immunohistological study by Andersson et al. the cells were classified into FcεRI positive/negative in a binary manner depending on the detection limit of the technique. We also observed a big variation of expression between individuals (Figure 3) and in line with our results this has previously been shown to be true also for human skin mast cells ^31^. The reason for the variation could be manifold as the surface expression of FcεRI can be regulated in many different ways. It is, for example, upregulated by IL-4 and stabilized on the cell surface by the binding of IgE antibodies^32^ and recently is was described that IL-33 downregulates the expression ^33,34^, indicating that the state of inflammation in the tissue could influence the FcεRI expression.

Human lung mast cells have been shown to be heterogenous, classically this have been studied using immunohistochemistry in a binary manner and they have been divided into MCT and MCTC, depending on whether or not the mast cell proteases chymase and CPA3 are detectable [6]. How this heterogeneity is reflected on heterogenous expression of cell surface markers is scarcely investigated. We investigated the heterogeneity of cell surface markers using flow cytometry in a quantitative manner and did not find any markers that distinctly and consistently divided the mast cells into subpopulations with a bi- or multimodal distribution (data not shown). We did however find several markers with considerable continuous variation of expression within the mast cell populations (Table 2) and co-staining revealed that six of these markers correlated, including SUSD2, CD49a, CD326, CD34, CD66 and HLA-DR (Figure 3). To investigate if these markers correlated to the classical mast cell subpopulations, MC_T_ and MC_TC_, we co-stained SUSD2 with CPA3, but no correlation was detected ruling out the possibility that they are extracellular markers of the classical mast cell subtypes (Figure 4G). CD344 did not correlate to MC_T_ and MC_TC_ profile either (data not shown). CD88 have been reported to be a cell surface marker that distinguish the MC_TC_ from the MC_T_ subset^35^, however in our hands we did not detect any expression of CD88 on the HLMC (Figure 2), thus we were unable to find an extracellular marker that distinguishes the classical mast cell subsets.

Considering that one of our six correlating heterogeneity markers, CD34, is expressed on circulating mast cell progenitors ^36^, we speculated that these markers could identify cells in different stages of maturation. However, if that was the case one would expect that cells with a high expression of CD34, and by correlation all the other heterogeneity markers, to be small and contain few granules similar to mast cell progenitors^36^. To the contrary, the cells had a higher FSC and SSC (Figure 4 H-J), suggesting that they are bigger and more granular, and therefore they are not likely to be immature mast cells. SUSD2 is a marker for pluripotent^20^ and mesenchymal^19^ stem cells but it is also expressed in certain cancers where it has been linked to proliferation^21^, thus one could imagine that cells high in SUSD2 are proliferating. However, we could not detect any staining of the proliferation marker Ki67 in the HLMC (Supplementary figure S2). We also saw varying expression of HLA-DR, an MHC class II receptor that presents antigens to CD4^+^ T-cells, suggesting that cells high in the heterogeneity markers could be able to present antigen and activate CD4^+^ T cells. There have initially been conflicting results from murine experiments regarding whether or not mast cells are able to present antigen and activate T cells via MHC II (reviewed in ^37^). However, human mast cells that are present in close proximity to T cells in tonsils express HLA-DR and CD80, indicating that they can present antigen to CD4^+^ T cells ^38^. *In vitro* derived human mast cells from CD34+ progenitors and *ex vivo* human skin mast cells have been shown to express MHC II and co-stimulatory ligands when stimulated with IFN-*γ* and activate T cells in an antigen-dependent manner ^38,39^. In this context it is worth noting that the co-stimulatory ligands for T cell activation, CD80, CD86, CTLA-4 (CD152), OX40L (CD252), Tim-1, Tim-4, 41BB-L (CD137L), ICOS-L (CD275), CD70, CD40, LIGHT (CD258) and CD112 were not detected on the HLMC, while CD48, CD58, CD155 and HVEM (CD270) were expressed (Figure 2). The co-inhibitory ligands PD-L1 (CD274) and PD-L2 (B7-DC, CD273) were also expressed, while Galectin-9 was not detected (Figure 2)^40^. Thus, the HLMCs are endowed with receptors/ligands given them possibilities to interact with and regulate T cells and the adaptive immunity ^41,42^.

In summary, we found expression of 102 cell surface antigens on HLMC, several of which had a high continuous variability of expression within the mast cell population. The expression of six of these markers correlated to each other (SUSD2, CD49a, CD326, CD34, CD66 and HLA-DR) and the size and granularity of the cells. Further studies are needed to determine how these cells differ functionally. To the contrary of the dogma of distinct mast cell subtypes, we demonstrate a continuous nature of HLMC heterogeneity.

## Acknowledgements

We thank Andrew Walls for the generous gift of CPA3 antibody. This study was supported by grants from the Swedish Research Council; the Heart-Lung Foundation; The Swedish Cancer Society; the Ellen, Walter and Lennart Hesselman’s foundation; Tore Nilssons Foundation; the Lars Hiertas memory fund; the Konsul Th C Burghs Foundation; the Tornspiran Foundation; the O. E. and Edla Johanssons Foundation; the Swedish Society for Medical Research; The ChAMP (Centre for Allergy Research Highlights Asthma Markers of Phenotype) consortium funded by the Swedish Foundation for Strategic Research; the AstraZeneca & Science for Life Laboratory Joint Research Collaboration; and the Karolinska Institutet.

## Notes

### Competing Interest Statement

The authors have declared no competing interest.

