## Supplementary table and figures for "Immunoprofiling reveals novel mast cell receptors and a continuous nature of human lung mast cell heterogeneity"

Supplementary Table S1. Antibodies included in the LEGENDScreen human cell Screening kit

| Specificity | Clone | Specificity | Clone |
|---|---|---|---|
| CD1a | HI149 | CD49f | GoH3 |
| CD1b | SN13 (K5- 1B8) | CD50 (ICAM-3) | CBR-IC3/1 |
| CD1c | L161 | CD51 | NKI-M9 |
| CD1d | 51.1 | CD51/61 | 23C6 |
| CD2 | RPA-2.10 | CD52 | HI186 |
| CD3 | HIT3a | CD53 | HI29 |
| CD4 | RPA-T4 | CD54 | HA58 |
| CD5 | UCHT2 | CD55 | JS11 |
| CD6 | BL-CD6 | CD56 (NCAM) | HCD56 |
| CD7 | CD7-6B7 | CD57 | HCD57 |
| CD8a | HIT8a | CD58 | TS2/9 |
| CD9 | HI9a | CD59 | p282 (H19) |
| CD10 | HI10a | CD61 | VI-PL2 |
| CD11a | HI111 | CD62E | HAE-1f |
| CD11b | ICRF44 | CD62L | DREG-56 |
| CD11b (activated) | CBRM1/5 | CD62P (P-Selectin) | AK4 |
| CD11c | 3.9 | CD63 | H5C6 |
| CD13 | WM15 | CD64 | 10.1 |
| CD14 | M5E2 | CD66a/c/e | ASL-32 |
| CD15 (SSEA-1) | W6D3 | CD66b | G10F5 |
| CD16 | 3G8 | CD69 | FN50 |
| CD18 | TS1/18 | CD70 | 113-16 |
| CD19 | HIB19 | CD71 | CY1G4 |
| CD20 | 2H7 | CD73 | AD2 |
| CD21 | Bu32 | CD74 | LN2 |
| CD22 | HIB22 | CD79b | CB3-1 |
| CD23 | EBVCS-5 | CD80 | 2D10 |
| CD24 | ML5 | CD81 | 5A6 |
| CD25 | BC96 | CD82 | ASL-24 |
| CD26 | BA5b | CD83 | HB15e |
| CD27 | O323 | CD84 | CD84.1.21 |
| CD28 | CD28.2 | CD85a (ILT5) | MKT5.1 |
| CD29 | TS2/16 | CD85d (ILT4) | 42D1 |
| CD30 | BY88 | CD85g (ILT7) | 17G10.2 |
| CD31 | WM59 | CD85h (ILT1) | 24 |
| CD32 | FUN-2 | CD85j (ILT2) | GHI/75 |
| CD33 | WM53 | CD85k (ILT3) | ZM4.1 |
| CD34 | 581 | CD86 | IT2.2 |
| CD35 | E11 | CD87 | VIM5 |
| CD36 | 5-271 | CD88 | S5/1 |
| CD38 | HIT2 | CD89 | A59 |
| CD39 | A1 | CD90 (Thy1) | 5E10 |
| CD40 | HB14 | CD93 | VIMD2 |
| CD41 | HIP8 | CD94 | DX22 |
| CD42b | HIP1 | CD95 | DX2 |
| CD43 | CD43-10G7 | CD96 | NK92.39 |
| CD44 | BJ18 | CD97 | VIM3b |
| CD45 | HI30 | CD99 | HCD99 |
| CD45RA | HI100 | CD100 | A8 |
| CD45RB | MEM-55 | CD101 (BB27) | BB27 |
| CD45RO | UCHL1 | CD102 | CBR-IC2/2 |
| CD46 | TRA-2-10 | CD103 | Ber-ACT8 |
| CD47 | CC2C6 | CD104 | 58XB4 |
| CD48 | BJ40 | CD105 | 43A3 |
| CD49a | TS2/7 | CD106 | STA |
| CD49c | ASC-1 | CD107a (LAMP-1) | H4A3 |
| CD49d | 9F10 | CD108 | MEM-150 |
| CD49e | NKI-SAM-1 | CD109 | W7C5 |

| Specificity | Clone |
|---|---|
| CD111 | R1.302 |
| CD112 (Nectin-2) | TX31 |
| CD114 | LMM741 |
| CD115 | 9-4D2-1E4 |
| CD116 | 4H1 |
| CD117 (c-kit) | 104D2 |
| CD119 | GIR-208 |
| CD122 | TU27 |
| CD123 | 6H6 |
| CD124 | G077F6 |
| CD126 (IL-6Rα) | UV4 |
| CD127 (IL-7Rα) | A019D5 |
| CD129 (IL-9 R) | AH9R7 |
| CD131 | 1C1 |
| CD132 | TUGh4 |
| CD134 | Ber-ACT35 (ACT35) |
| CD135 | BV10A4H2 |
| CD137 (4-1BB) | 4B4-1 |
| CD137L (4-1BB Ligand) | 5F4 |
| CD138 | DL-101 |
| CD140a | 16A1 |
| CD140b | 18A2 |
| CD141 | M80 |
| CD143 | 5-369 |
| CD144 | BV9 |
| CD146 | SHM-57 |
| CD148 | A3 |
| CD150 (SLAM) | A12 (7D4) |
| CD152 | L3D10 |
| CD154 | 24-31 |
| CD155 (PVR) | SKII.4 |
| CD156c (ADAM10) | SHM14 |
| CD158a/h | HP-MA4 |
| CD158b | DX27 |
| CD158d | mAb 33 (33) |
| CD158e1 | DX9 |
| CD158f | UP-R1 |
| CD161 | HP-3G10 |
| CD162 | KPL-1 |
| CD163 | GHI/61 |
| CD164 | 67D2 |
| CD165 | SN2 (N6- D11) |
| CD166 | 3A6 |
| CD167a (DDR1) | 51D6 |
| CD169 | 7-239 |
| CD170 (Siglec-5) | 1A5 |
| CD172a (SIRPa) | SE5A5 |
| CD172b (SIRPb) | B4B6 |
| CD172g (SIRPg) | LSB2.20 |
| CD178 (Fas-L) | NOK-1 |
| CD179a | HSL96 |
| CD179b | HSL11 |
| CD180 (RP105) | MHR73-11 |
| CD181 (CXCR1) | 8F1/CXCR1 |
| CD182 (CXCR2) | 5E8/CXCR2 |
| CD183 | G025H7 |
| CD184 (CXCR4) | 12G5 |

| Specificity | Clone |
|---|---|
| CD193 (CCR3) | 5E8 |
| CD195 (CCR5) | T21/8 |
| CD196 | G034E3 |
| CD197 (CCR7) | G043H7 |
| CD200 (OX2) | OX-104 |
| CD200 R | OX-108 |
| CD201 (EPCR) | RCR-401 |
| CD202b (Tie2/Tek) | 33.1 (Ab33) |
| CD203c (E-NPP3) | NP4D6 |
| CD205 (DEC- 205) | HD30 |
| CD206 (MMR) | 15-2 |
| CD207 (Langerin) | 10E2 |
| CD209 (DC-SIGN) | 9E9A8 |
| CD210 (IL- 10 R) | 3F9 |
| CD213a2 | SHM38 |
| CD215 (IL- 15Rα) | JM7A4 |
| CD218a (IL-18Rα) | H44 |
| CD220 | B6.220 |
| CD221 (IGF-1R) | 1H7/CD221 |
| CD226 (DNAM-1) | 11A8 |
| CD229 (Ly-9) | HLy-9.1.25 |
| CD231 (TALLA) | SN1a (M3- 3D9) |
| CD235ab | HIR2 |
| CD243 | UIC2 |
| CD244 (2B4) | C1.7 |
| CD245 (p220/240) | DY12 |
| CD252 (OX40L) | 11C3.1 |
| CD253 (Trail) | RIK-2 |
| CD254 | MIH24 |
| CD255 (TWEAK) | CARL-1 |
| CD257 (BAFF, BLYS) | T7-241 |
| CD258 (LIGHT) | T5-39 |
| CD261 | DJR1 |
| CD262 | DJR2-4 (7-8) |
| CD263 | DJR3 |
| CD266 | ITEM-1 |
| CD267 (TACI) | 1A1 |
| CD268 (BAFF-R) | 11C1 |
| CD270 (HVEM) | 122 |
| CD271 | ME20.4 |
| CD273 (B7- DC, PD-L2) | 24F.10C12 |
| CD274 (B7- H1, PD-L1) | 29E.2A3 |
| CD275 (B7- H2) | 9F.8A4 |
| CD276 | MIH42 |
| CD277 | BT3.1 |
| CD278 (ICOS) | C398.4A |
| CD279 (PD-1) | EH12.2H7 |
| CD282 (TLR2) | TL2.1 |
| CD284 (TLR4) | HTA125 |
| CD286 (TLR6) | TLR6.127 |
| CD290 | 3C10C5 |
| CD294 | BM16 |
| CD298 | LNH-94 |
| CD300e (IREM-2) | UP-H2 |
| CD300F | UP-D2 |
| CD301 | H037G3 |
| CD303 | 201A |

| Specificity | Clone |
|---|---|
| CD304 | 12C2 |
| CD307 | 509f6 |
| CD307d (FcRL4) | 413D12 |
| CD314 (NKG2D) | 1D11 |
| CD317 | RS38E |
| CD318 (CDCP1) | CUB1 |
| CD319 (CRACC) | 162.1 |
| CD324 (E-Cadherin) | 67A4 |
| CD325 | 8C11 |
| CD326 (Ep- CAM) | 9C4 |
| CD328 (Siglec-7) | 6-434 |
| CD334 (FGFR4) | 4FR6D3 |
| CD335 (NKp46) | 9E2 |
| CD336 (NKp44) | P44-8 |
| CD337 (NKp30) | P30-15 |
| CD338 (ABCG2) | 5D3 |
| CD340 (erbB2/ HER-2) | 24D2 |
| CD344 | CH3A4A7 |
| CD351 | TX61 |
| CD352 (NTB-A) | NT-7 |
| CD354 (TREM-1) | TREM-26 |
| CD355 (CRTAM) | Cr24.1 |
| CD357 (GITR) | 621 |
| CD360 (IL- 21R) | 2G1-K12 |
| β2- microglobulin | 2M2 |
| BTLA | MIH26 |
| C3AR | hC3aRZ8 |
| C5L2 | 1D9-M12 |
| CCR10 | 6588-5 |
| CLEC12A | 50C1 |
| CLEC9A | 8F9 |
| CX3CR1 | 2A9-1 |
| CXCR7 | 8F11-M16 |
| OPRD | DOR7D2A4 |
| DLL1 | MHD1-314 |
| DLL4 | MHD4-46 |
| DR3 (TRAMP) | JD3 |
| EGFR | AY13 |
| erbB3/HER-3 | 1B4C3 |
| FcεRIα | AER-37 (CRA-1) |
| FcRL6 | 2H3 |
| Galectin-9 | 9M1-3 |
| GARP (LRRC32) | 7B11 |
| HLA-A,B,C | W6/32 |
| HLA-A2 | BB7.2 |
| HLA-DQ | HLADQ1 |
| HLA-DR | L243 |
| HLA-E | 3D12 |
| HLA-G | 87G |
| IFNGR2 | 2HUB-159 |
| Ig light chain k | MHK-49 |
| Ig light chain λ | MHL-38 |
| IgD | IA6-2 |
| IgM | MHM-88 |
| IL-28RA | MHLICR2a |
| Integrin α9β1 | Y9A2 |

| Specificity | Clone |
|---|---|
| integrin β5 | AST-3T |
| integrin β7 | FIB504 |
| Jagged 2 | MHJ2-523 |
| LAP | TW4-6H10 |
| LT-bR | 31G4D8 |
| Mac-2 (Galectin-3) | Gal397 |
| MAIR-II | TX45 |
| MICA/MICB | 6D4 |
| MSC (W3D5) | W3D5 |
| MSC (W5C5) | W5C5 |
| MSC (W7C6) | W7C6 |
| MSC and NPC (W4A5) | W4A5 |
| MSCA-1 (MSC, W8B2) | W8B2 |
| NKp80 | 5D12 |
| Notch 1 | MHN1-519 |
| Notch 2 | MHN2-25 |
| Notch 3 | MHN3-21 |
| Notch 4 | MHN4-2 |
| NPC (57D2) | 57D2 |
| Podoplanin | NC-08 |
| Pre-BCR | HSL2 |
| PSMA | LNI-17 |
| Siglec-10 | 5G6 |
| Siglec-8 | 7C9 |
| Siglec-9 | K8 |
| SSEA-1 | MC-480 |
| SSEA-3 | MC-631 |
| SSEA-4 | MC-813-70 |
| SSEA-5 | 8E11 |
| TCR g/d | B1 |
| TCR Vβ13.2 | H132 |
| TCR Vβ23 | αHUT7 |
| TCR Vβ8 | JR2 (JR.2) |
| TCR Vβ9 | MKB1 |
| TCR Vδ2 | B6 |
| TCR Vg9 | B3 |
| TCR Vα24- Jα18 | 6B11 |
| TCR Vα7.2 | 3C10 |
| TCR α/β | IP26 |
| Tim-1 | 1D12 |
| Tim-3 | F38-2E2 |
| Tim-4 | 9F4 |
| TLT-2 | MIH61 |
| TRA-1-60-R | TRA-1-60-R |
| TRA-1-81 | TRA-1-81 |
| TSLPR (TSLP-R) | 1B4 |
| Ms IgG1, κ ITCL | MOPC-21 |
| Ms IgG2a, κ ITCL | MOPC-173 |
| Ms IgG2b, κ ITCL | MPC-11 |
| Ms IgG3, κ ITCL | MG3-35 |
| Ms IgM, κ ITCL | MM-30 |
| Rat IgG1, κ ITCL | RTK2071 |
| Rat IgG2a, κ ITCL | RTK2758 |
| Rat IgG2b, κ ITCL | RTK4530 |
| Rat IgM, κ ITCL | RTK2118 |
| AH IgG, ITCL | HTK888 |

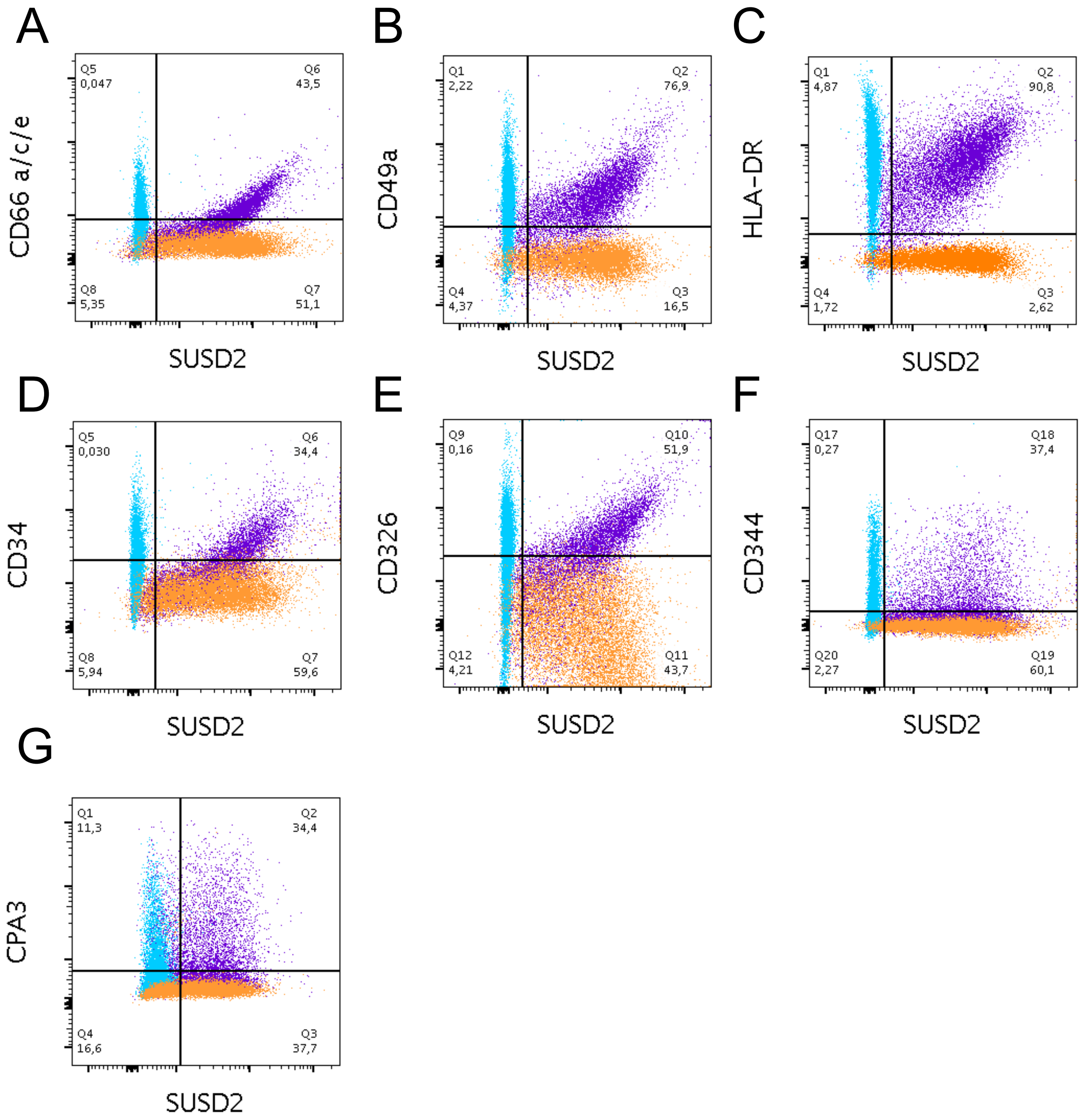


**Supplementary Figure S1. Correlations between markers with high CV including FMO controls**

HLMCs were co-stained with SUSD2 (A-H), CD66a/c/e (A), CD49a (B), HLA-DR (C), CD34 (D), CD326 (E), CD344 (F), and CPA3 (G). Co-stainings with SUSD2 is shown in purple, FMO control for SUSD2 in blue, and FMO controls for all other markers in orange. Representatives of 4 donors are shown.

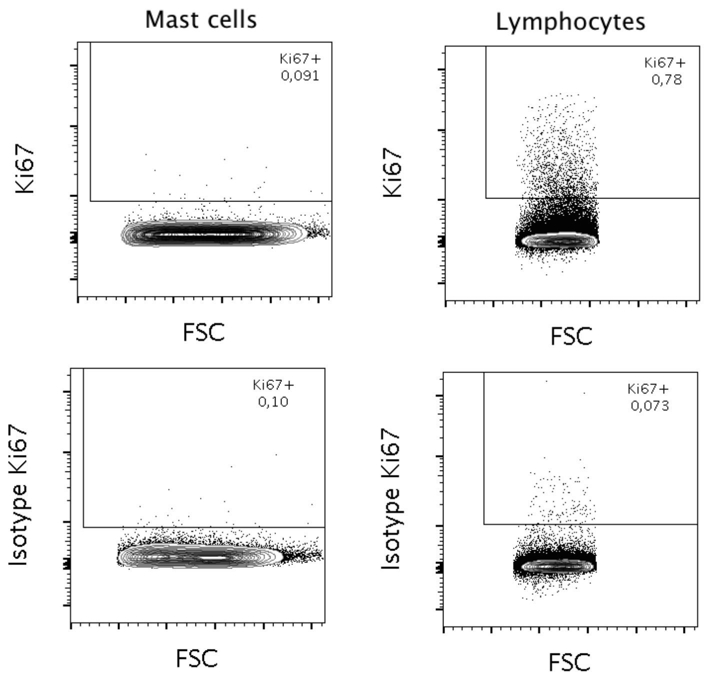


**Supplementary Figure S2. No proliferation detected in HLMCs**.

HLMCs gated as $CD45^{+}$, $CD14^{low}$ and $CD117^{high}$ were stained intracellularly with the proliferation marker Ki67. As a positive control for the Ki67 staining, cells enriched with lymphocytes were also gated as $SSC^{low}FSC^{low}CD45^{+}$.
